# Modular modeling improves the predictions of genetic variant effects on splicing

**DOI:** 10.1101/438986

**Authors:** Jun Cheng, Thi Yen Duong Nguyen, Kamil J Cygan, Muhammed Hasan Çelik, William G Fairbrother, Žiga Avsec, Julien Gagneur

## Abstract

Predicting the effects of genetic variants on splicing is highly relevant for human genetics. We describe the framework MMSplice (modular modeling of splicing) with which we built the winning model of the CAGI 2018 exon skipping prediction challenge. The MMSplice modules are neural networks scoring exon, intron, and splice sites, trained on distinct large-scale genomics datasets. These modules are combined to predict effects of variants on exon skipping, alternative donor and acceptor sites, splicing efficiency, and pathogenicity, with matched or higher performance than state-of-the-art. Our models, available in the repository Kipoi, apply to variants including indels directly from VCF files.

## Introduction

Genetic variants altering splicing constitute one of the most important class of genetic determinants of rare [1, 2] and common [2] diseases. However, the accurate prediction of variant effect on splicing remains challenging.

Splicing is the outcome of multiple processes. It is a two-step catalytic process in which a donor site is first attacked by an intronic adenosine to form a branch-point. In a second step, the acceptor site is cleaved and spliced (i.e. joined) to the 3’end of the donor site. The sequences of the donor site, of the acceptor site, and of the intronic region surrounding the branchpoint, which are recognized during spliceosome assembly contribute to splicing regulation [3]. Moreover, many regulatory elements such as exonic splicing enhancers (ESEs) and silencers (ESSs), intronic splicing enhancers (ISEs) and silencers (ISSs) also play key regulatory roles (reviewed by [4]). In addition to genetic variants at splice consensus sequence, distal elements can also affect splicing and cause disease [5]. Hence, predictive models of splicing need to integrate these various types of sequence elements.

Previous human splice variant interpretation methods can be grouped into two categories.

One category consists of algorithms that score sequence for being bona fide splice regulatory elements including the splice sites [6, 7], the exonic and intronic enhancers and silencers [8, 9, 10, 11, 12, 13]. Variants can be scored with respect to these regulatory elements by comparing predictions for the reference sequence and the alternative sequence containing the genetic variant of interest. However, although methods combining several of these scores have been proposed, including Human Splicing Finder [14], MutPred splice [15], and more recently SPiCE [16], the resulting physical and quantitative effect of these variants on splicing remains difficult to assess with these algorithms.

The second category of models aimed at predicting the effect of genetic variants on relative amounts of alternative splicing isoforms quantitatively [17, 18, 19]. In this context, a quantitative measure that has retained much attention in the literature is the percent spliced-in (PSI, also denoted Ψ), which quantifies exon skipping. Ψ is defined as the fraction of transcripts that contains a given exon [20]. It can be estimated as the fraction of exon-exon junction reads from an RNA-seq sample supporting inclusion of an exon of interest, over the sum of these reads plus those supporting the exclusion of this exon [20]. Two state-of-the-art models for predicting Ψ from sequence are SPANR [17] and HAL [18]. A related quantity, Ψ_5_ quantifies for a given donor site the fraction of spliced transcripts with a particular alternative 3’splice site (A3SS). The quantity Ψ_3_ has been analogously defined to quantify alternative 5’splice sites (A5SS) [21]. The recently published algorithm COSSMO [19] predicts Ψ_5_ from sequence by modeling a competition between alternative acceptor sites for a given donor site, and analogously for Ψ_3_. COSSMO has shown superior performance over MaxEntScan[7] on predicting the most frequently used splice site among competing ones. Furthermore, splicing efficiency has been proposed to quantify the amount of precursor RNA that undergo splicing (exon-skipped or misspliced transcripts are ignored) at a given splice site by comparing the amount of RNA-seq reads spanning an exon-intron boundary of interest to the corresponding exon-exon junction reads [22]. The latest model to predict variant effect on splicing efficiency is the SMS score, which is based on scores for exonic 7-mers estimated from a recently published saturation mutagenesis assay [23]. However, no model can be applied to all the above mentioned splicing quantities, although they are influenced by common regulatory elements. Furthermore, none of these software handle variant calling format (VCF) files natively, making their integration into genetic diagnostics pipelines cumbersome. Also, these software often do not handle indels (insertions and deletions), although indels are potentially the most deleterious variants.

Here we trained building-block modules separately for the exon, the acceptor site, the donor site and for intronic sequence close to the donor and close to the acceptor sites. This modular approach allowed leveraging rich datasets from two high-throughput perturbation assays focusing on distinct aspects of splicing: i) a massive parallel reporter assay (MPRA) with millions of random short sequences in intron or exon probe splicing regulatory sequences in great depth [18], and ii) a high-throughput quantify effect of nature occurring variants from all splicing regulatory regions on splicing of their harbouring exon [24]. These building-block modules could then be combined into distinct models predicting effects of variants on Ψ, Ψ_5_, Ψ_3_, splicing efficiency, and one model predicting splice variant pathogenicity trained on the database ClinVar [25]. We outperform state-of-the-art-models for each task but Ψ_3_, on which MMSplice and HAL both are the best. In particular, our model of exon-skipping ranked first at the 2018 CAGI challenge https://genomeinterpretation.org. All our models are available open source in the model zoo Kipoi [26] and can be applied for variant effect prediction directly from VCF files.

## Results

### Modular modeling strategy

We designed neural networks to score five potentially overlapping splicing-relevant sequence regions: the donor site, the acceptor site, the exon, as well as the 5’end and the 3’end of the intron (Fig. 1A). The donor and the acceptor models were trained to predict annotated intron-exon and exon-intron boundaries from GENCODE 24 genome annotation (Fig. 1A). The exon and intron models were trained from a MPRA that probed the effect of millions of random sequences altering either the exonic 3’end and the intronic 5’end for alternative 5’splicing (A5SS, quantified by Ψ_3_), or the exonic 5’end and the intronic 3’end for alternative 3’splicing (A3SS, quantified by Ψ_5_) (Fig. 1A) [18]. For later use, the modules were defined as the corresponding neural network models without the last activation layer. We have two intron modules, the intron 5’module that scores intron from the donor side and the intron 3’module that scores intron from the acceptor side. Likewise we have two exon modules, the exon 5’module that trained from A3SS and exon 3’module that trained from A5SS (Supplement Fig. S1). To score exonic sequence, only one of the exonic module is applied depending on the alternative splicing quantity. Training data and module architecture are summarized in Table 1. Next, we combined these modules to predict how genetic variants lead to i) differences in Ψ, ii) differences Ψ_3_, iii) differences in Ψ_5_, iv) differences in splicing efficiency and v) to disease or benign phenotypes according to the ClinVar database (Fig. 1B). Specifically, we trained one linear model on top of the modules to predict ΔΨ. The same linear model was applied to predict ΔΨ_5_ and ΔΨ_3_ by modeling the competition of two alternative exons. Another linear model was trained to predict change of splicing efficiency and a logistic regression model was trained to predict variant pathogenicity from the modules (Fig. 1B).

**Figure 1.**
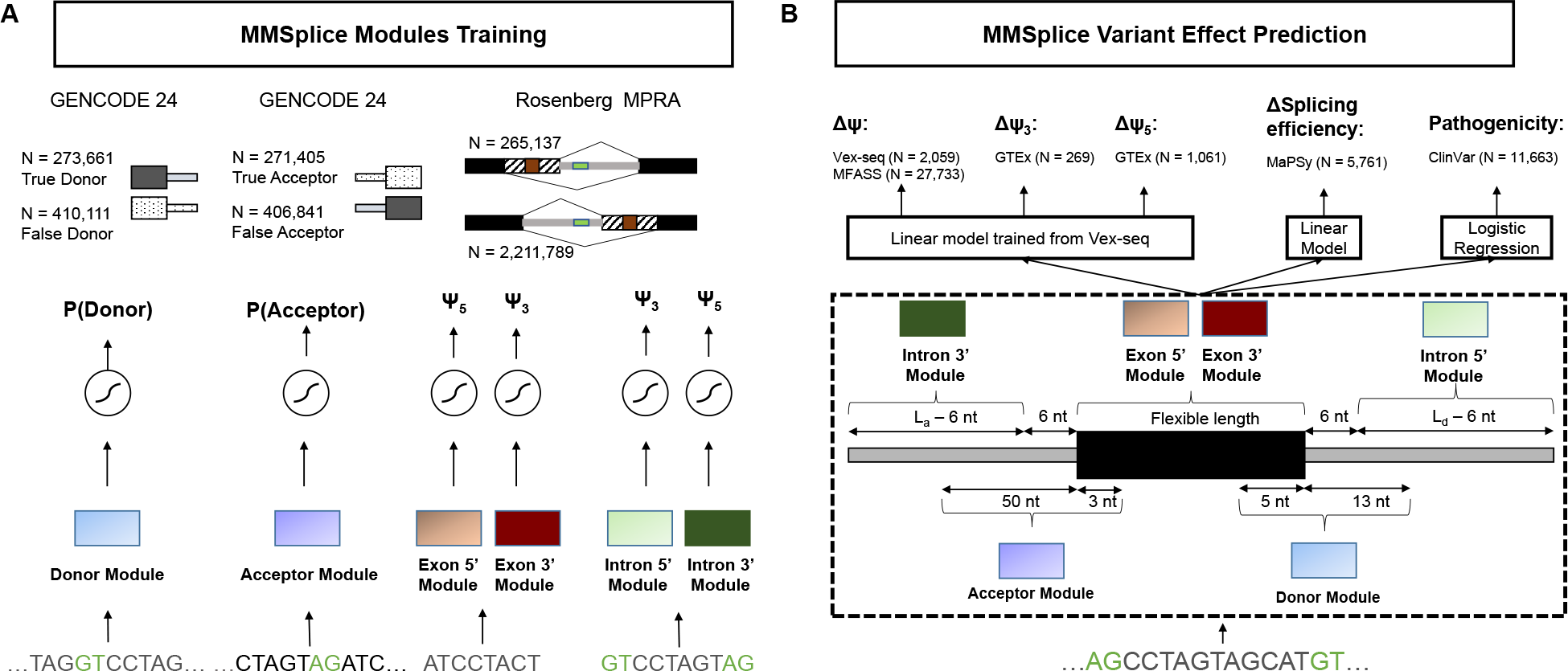
Individual modules of MMSplice and their combination to predict the effect of genetic variants on various splicing quantities. **(A)** MMSplice consists of six modules scoring sequences from donor, acceptor, exon and intron sites. Modules were trained with rich genomics dataset probing the corresponding regulatory regions. **(B)** Modules from (A) are combined with a linear model to score variant effect on exon skipping (ΔΨ), alternative donor (ΔΨ_3_) or alternative acceptor site (ΔΨ_5_), splicing efficiency, and they are combined with a logistic regression model to predict variant pathogenicity. *L_a_* and *L_d_* stands for the length of intron sequence taken from the acceptor and donor side.

**Table 1.** Summary of trained modules and their training data.

| Module | Training Data | Strategy |
| --- | --- | --- |
| Intron 3'module | MPRA[18] intronic random sequence | Share conv layer, dense layer specific for 3'intronic random sequence (A3SS library) |
| Acceptor module | GENCODE 24 annotation | Classify annotated acceptors versus random sequence |
| Exon 5'module | MPRA[18] exonic random sequence | Share conv layer, dense layer specific for 5'exonic random sequence (A3SS library) |
| Exon 3'module | MPRA[18] exonic random sequence | Share conv layer, dense layer specific for 3'exonic random sequence (A5SS library) |
| Donor module | GENCODE 24 annotation | Classify annotated donors and random sequence |
| Intron 5'module | MPRA[18] intronic random sequence | Share conv layer, dense layer specific for 5'intronic random sequence (A5SS library) |

### MMSplice improves the prediction of variant effect on exon skipping

To assess the performance of MMSplice for predicting causal effects of variants on exon skipping, we first considered the Vex-seq dataset [24]. Vex-seq is a high-throughput reporter assay that compared Ψ for constructs containing a reference sequence to Ψ for matching constructs containing one of 2,059 Exome Aggregation Consortium (ExAC [27]) variants. The difference of Ψ for the variant allele to the reference allele is denoted ΔΨ. These variants consisted of both single nucleotide variants as well as indels from exons and introns (20 nt upstream, 50 nt downstream). The data for the HepG2 cell line was accessed through the Critical Assessment of Genome Interpretation (CAGI) competition [28]. The 957 variants from chromosome 1 to chromosome 8 were provided as training data. The remaining 1,054 variants from chromosome 9 to 22 and chromosome X were held out for testing by the CAGI competition organizers and were not available throughout the development of the model. The test data consisted of 572 exonic and 526 intronic variants, and included 44 indels.

The Vex-seq experiment is an exon skipping assay, whereas our exon modules were trained for A5SS (Ψ_3_) and A3SS (Ψ_5_). Because of high redundancy between these two modules, we used the 5’module exon module as it was better at predicting exon skipping exonic variants on Vex-seq training data than the exon 3’module (*R* = 0.5 v.s *R* = 0.24, *P* = 0.001, bootstrap, Supplementary Fig. S2).

We built an MMSplice predictor for ΔΨ by training a linear model to combine the modular predictions and interaction terms between modules with overlapping scored regions from the Vex-seq training data (Methods, Eq. 2). We compared MMSplice with three state-of-the-art splicing variant scoring models: SPANR [17], HAL [18] and MaxEntScan [7] on the held-out Vex-seq test data. The methods HAL [18] and SPANR [17] have been reported to be the two best performed existing methods on a recent large-scale perturbation assay probing 27,733 rare variants [29], while MaxEntScan [7] was considered as a baseline reference model. SPANR scores exonic and intronic SNVs up to 300 nt around splice junctions. HAL scores exonic and donor (6 nt to the intron) variants. MaxEntScan scores [−3, +6] nt around the donor and [−20, +3] nt around the acceptor sites. The Vex-seq data was processed the same way for these models (Methods). Unlike the other methods SPANR does not take custom input sequences and could therefore score single nucleotide variants but not for indels.

On the Vex-seq data, MMSplice showed a large improvement over HAL and SPANR. First, MMSplice could score all 1,098 variants of the test set whereas HAL could only score 572 (52.1%) and SPANR 966 (88%) of them. Second, the difference in Ψ predicted by MMSplice correlated better globally but also when restricted to the respective variants scored by the other methods (*R* = 0.68 for MMSplice v.s. *R* = 0.41, 0.24 for HAL and SPANR respectively, both comparison *P* = 0.001, bootstrap, Fig. 2). A higher performance than other models was also obtained even when we bluntly summed the prediction scores from the 5 modules without fitting any parameter to the Vex-seq training data (*R* = 0.67, and *R* = 0.66 when using the exon 3’module in place of the exon 5’module, Supplement Fig. S3). This shows that the superior performance of our model is primarily due to the modules not the combination linear model that was trained from Vex-seq training data.

**Figure 2.**
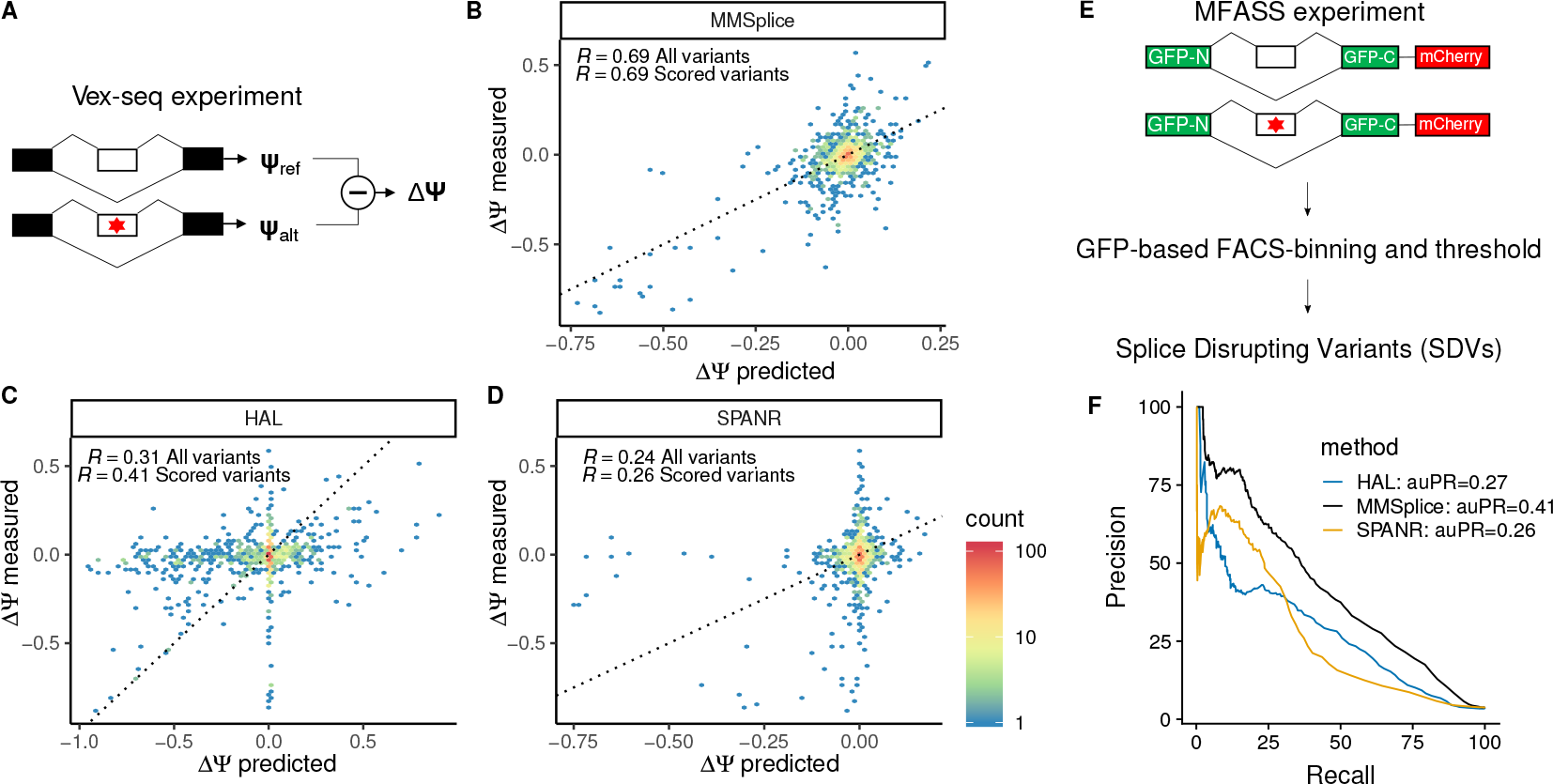
MMSplice improves the prediction of variant effect on exon skipping. **(A)** Schema of the Vex-seq experiment [24]. The effect of 2,059 ExAC variants (red star) from or adjacent to 110 alternative exons were tested with reporter genes by measuring percent splice-in of the reference sequence (Ψ_ref_) and of the alternative (Ψ_alt_) by RNAseq. **(B-D)** Measured (y-axis) versus predicted (x-axis) Ψ differences between alternative and reference sequence for MMSplice (B), HAL[18] (C), and SPANR[17] (D) on Vex-seq test data. Color scale represents counts in hexagonal bins. The black line marks the y=x diagonal. Pearson correlations (*R*) were calculated i) for all variants and ii) only for the variants the considered model can score. **(E)** Schema of MFASS experiment [29]. Exon skipping effects of 27,733 ExAC SNVs (red star) spanning or adjacent to 2,339 exons were tested by genome integration of designed construct. Splice-disrupting variant (SDV) is defined as a variant that change an exon with original exon inclusion index ≥ 0.5 by at least 0.5. **(F)** Precision-recall curve of MFASS SDVs classification based on model predicted ΔΨ. Precision-recall curve for all three models were calculated for the sets of variants they can score. MMSplice (black) scored all 27,733 variants, SPANR (yellow) scored 27,663 variants (1,048 SDVs), and HAL (blue) scored 14,353 variants (489 SDVs).

We further compared our prediction for donor and acceptor site variants with the popular model MaxEntScan [7]. MMSplice performed better both in donor sequence (*R* = 0.87 for MMSplice v.s. 0.66 for MaxEntSan5, *P* = 0.001, bootstrap, Supplementary Fig. S4) and acceptor sequence (*R* = 0.81 for MMSplice v.s. 0.69 for MaxEntSan3, *P* = 0.001, bootstrap, Supplementary Fig. S5), when restricted to the subset of variants that MaxEntScan3 could score (42 donor variants and 149 acceptor variants). HAL performed better (*R* = 0.71) than MaxEntScan5 (*R* = 0.66) but worse than MMSplice (*R* = 0.87) on donor variants (*P* = 0.001 for both comparisons, bootstrap, Supplementary Fig. S4).

Altogether, this benchmark on large scale perturbation experiment demonstrates that MMSplice outperforms SPANR, HAL and MaxEntScan on predicting causal effects of genetic variants on exon skipping, by covering more variants and also by providing more accurate predictions. Our model also ranked the first in the 2018 CAGI Vex-seq competition. A joint publication with the organizers and challengers is in the planning.

### MMSplice classifies rare splice disrupting variants with higher precision and recall

To further compare models on predicting exon skipping level with independent datasets that no model has been trained on, we used the splicing functional assay from Cheung et al [29]. Cheung et al found 1,050 splice-disrupting variants (SDVs), the majority are extremely rare, after examining 27,733 ExAC single-nucleotide variants (SNV) with Multiplexed Functional Assay of Splicing using Sort-seq (MFASS) (Fig. 2E). The author benchmarked several variant effect prediction methods including conservation based methods like CADD [30], phastCons [31] and the state-of-the-art splicing variant scoring tools HAL and SPANR. Among all, the two splicing variant scoring methods performed much better than the others, thus MMSplice was compared with those two. MMSplice model with the final combination linear model trained from Vex-seq training data was applied to classify SDVs based on predicted ΔΨ solely from sequence. Our model achieved overall higher auPR (MMSplice 0.41, HAL 0.27, SPANR 0.26, *P* = 0.001 for both MMSplice v.s. HAL and MMSplice v.s. SPANR, bootstrap) when all models considering only their scored variants (Fig. 2F). In total, MMSplice scored all variants, SPANR scored 99.7% of all variants, while HAL scored only 51.8% of them. When considering exonic variants only, MMSplice (auPR=0.29) performed similar to HAL (auPR=0.27) (*P* = 0.326, bootstrap, Supplementary Fig. S6). For intronic variants, MMSplice had an auPR of 0.55 in comparison to 0.43 for SPANR (*P* = 0.001, bootstrap, Supplementary Fig. S6).

Overall, MMSplice demonstrated a substantital improvement over SPANR for both intronic and exonic variants and showed a similar performance than HAL for classifying exonic SDVs. This result also demonstrates the power of our model to score the effect of rare variants, for which association studies often lack power.

### MMSplice predicts variant effect on competing splice site selection with high accuracy

The MMSplice modular framework allows modeling alternative splicing events other than exon skipping. To demonstrate this and assess the performance of MMSplice on other alternative splicing events, we built MMSplice models to predict effect of variants around alternative donors on alternative 5’ splicing (A5SS, Ψ_3_) and variants around alternative acceptors on alternative 3’ splicing (A3SS) (Methods). Ψ_5_ and Ψ_3_ values for homozygous reference variants as well as with heterozygous and homozygous alternative variants were calculated from RNA-seq data of the GTEx consortium [32] (Methods). Here too, our MMSplice models allowed handling indels. One example is the insertion variant rs11382548 (chr11:61165731:C-CA). It is a splice site variant that turns a CG acceptor to an AG acceptor. It showed the largest ΔΨ_5_ among all assessed variants.

We benchmarked MMSplice against MaxEntScan, HAL and COSSMO. Overall, MMSplice (*R* = 0.66) significantly outperformed COSSMO (*R* = 0.5) (*P* = 0.016, bootstrap) and MaxEntScan (*R* = 0.46) (*P* = 0.001, bootstrap) and tie to HAL (*R* = 0.67, *P* = 0.558, bootstrap) on predicting ΔΨ_3_ (Fig. 3A-D). On predicting ΔΨ_5_, MMSplice (*R* = 0.57) again significantly outperformed both COSSMO (*R* = 0.37) and MaxEntScan (*R* = 0.44) (all *P* = 0.001, Fig. 3E-G). Even though HAL can predict A5SS donor variants well, the model has been trained for predicting A5SS and may not generalize well to other alternative splicing types. It only showed moderate performance when predicting donor variants from Vex-seq skipped exons (Supplement Fig. S4). In contrast, MMSplice showed consistent high performance across different types alternative splicing events.

**Figure 3.**
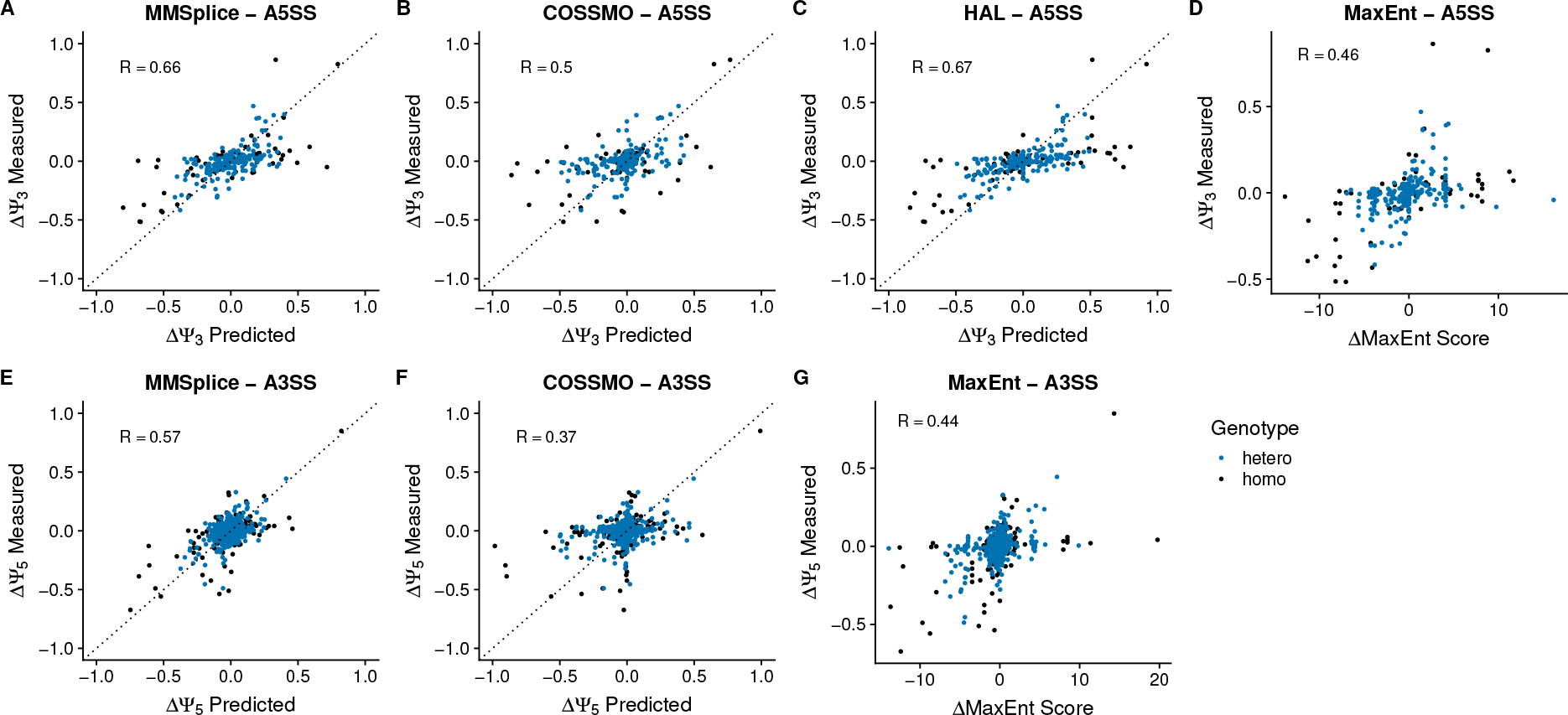
Evaluation of models predicting ΔΨ_5_ and ΔΨ_3_ on the GTEx dataset. GTEx variants around alternative spliced donors (3 nt in the exon and 6 nt in the intron) and acceptors (3 nt in the exon and 20 nt in the intron) were considered. Ψ_5_ (or Ψ_3_) of homozygous and heterozygous alternative variants as well as homozygous reference variants were calculated by taking the mean Ψ_5_ (or Ψ_3_) across individuals with the same genotype (excluding individuals with multiple variants within 300 nt around splice sites) on brain and skin (not sun exposed) samples. For donor variants, MMSplice (**A**) was benchmarked against COSSMO (**B**), HAL (**C**) and MaxEntScan (**D**). For acceptor variants, MMSplice (**E**) was benchmarked against COSSMO (**F**) and MaxEntScan (**G**).

MMSplice outperformed COSSMO for both donor and acceptor variants even though COSSMO was trained from estimated Ψ_5_ and Ψ_3_ values from GTEx data, while the training of MMSplice was independent from GTEx. MaxEntScan had similar performance as COSSMO for predicting ΔΨ_3_ (*R* = 0.46) and ΔΨ_5_ (*R* = 0.44)(Fig. 3).

### Prediction of splicing efficiency

We next used our modular approach to derive a model that predicts splicing efficiency, i.e. the proportion of spliced RNAs among spliced and unspliced RNAs [22]. We have done so in the context of a second 2018 CAGI challenge (Fig. 4A), whose training dataset is based on a massively parallel splicing assay (MaPSy [22]) and is described in Methods. This MaPSy dataset consists of splicing efficiencies 5,761 pairs of matched wild-type and mutated constructs, where each mutated construct differed from its matched wild-type by one exonic non-synonymous single-nucleotide variant (Methods). The assay has been done both with an *in vitro* splicing assay and *in vivo* by transfection into HEK293 cells (Methods). A test set of 797 construct pairs was held-out during the development of the model.

**Figure 4.**
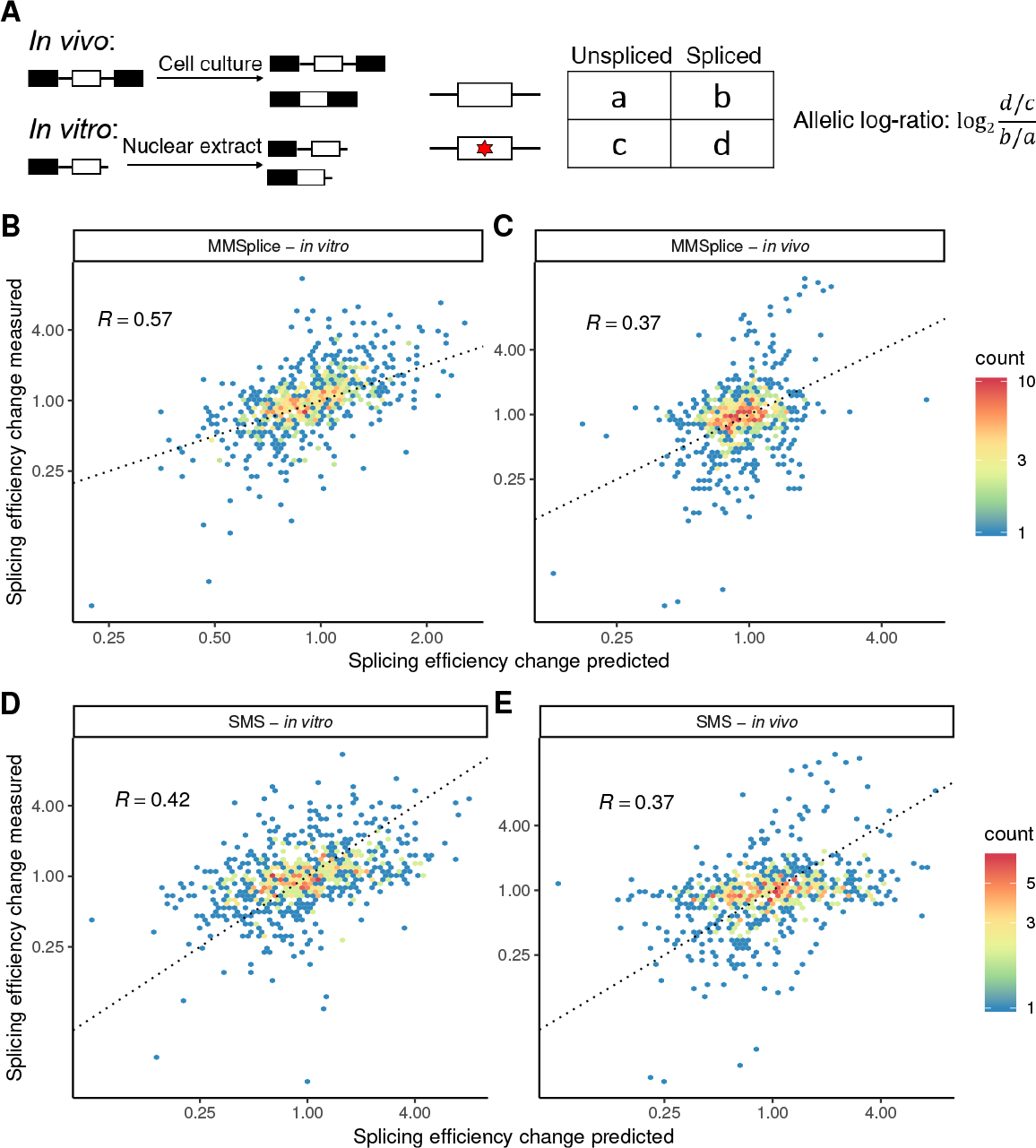
Splicing efficiency prediction. **(A)** MaPSy experiment (Methods). Effect of 5,761 published disease-causing exonic mutations on splicing efficiency are measured both *in vivo* and *in vitro*. Changes of splicing efficiency were quantified by allelic log-ratio. **(B-E)** Measured (y-axis) versus predicted (x-axis) allelic ratio for 797 variants in the test set for MMSplice (B-C) and the SMS score [23] (D-E). Dashed lines represent y=x line.

We trained a linear model on top of the modular predictions with MaPSy training data to predict differential splicing efficiency reported by the MaPSy data (Methods). This linear model was trained the same way as for Vex-seq except that the response was the allelic log-ratio (Fig. 4A and Methods) instead of Δ*logit*(Ψ). Our MMSplice model for differential splicing efficiencies predicted the effect of those non-synonymous mutations on the held-out test set reasonably well *in vitro* (*R* = 0.57, 4A) and well *in vivo* (*R* = 0.37, 4C). Also, our MMSplice model for differential splicing efficiencies outperformed the SMS score algorithm [23] on *in vitro* data (*P* = 0.001, bootstrap, 4D) and reached similar performance on the *in vivo* data (*P* = 0.524, bootstrap, 4E). Because splicing efficiency is defined as the ratio of spliced over precursor RNA *in vivo*, it is actually influenced both by variations in splicing but also by variations in RNA stability. Hence, our model may show a higher accuracy for the *in vitro* assay, because it involves only splicing-related factors, whereas degradation factors are also involved *in vivo*.

### MMSplice improves the prediction of splice variant pathogenicity

Predicting variant pathogenicity is a central task of genetic diagnosis. However, large amount of variants are annotated as “variant of uncertain significance” (VUS). A good splice variant effect prediction model can help interpreting VUSs. To evaluate the performance of MMSplice on predicting variant pathogenicity, we considered the ClinVar variants (version 20180429, [25]) that lie between 40 nt 5’ and 10 nt 3’ of an acceptor site or 10 nt either side of a donor site of a protein coding gene (Ensembl GRCh37 v75 annotation, Methods) as potentially affecting splicing. Among these variants, we aimed at discriminating between the 6,310 variants classified as pathogenic and the 4,405 variants classified as benign. To this end, we built an MMSplice model that implements a logistic regression on top of the MMSplice modules (Methods). Variants can potentially be in the vicinity of multiple exons. MMSplice handles this many-to-many relationship (Fig. 5A). Conveniently, MMSplice can be applied to a variant file in the standard format VCF [33] and a genome annotation file in the standard GTF format. Moreover, MMSplice is available as a Variant Effect Predictor Plugin (VEP, [34]) plugin.

**Figure 5.**
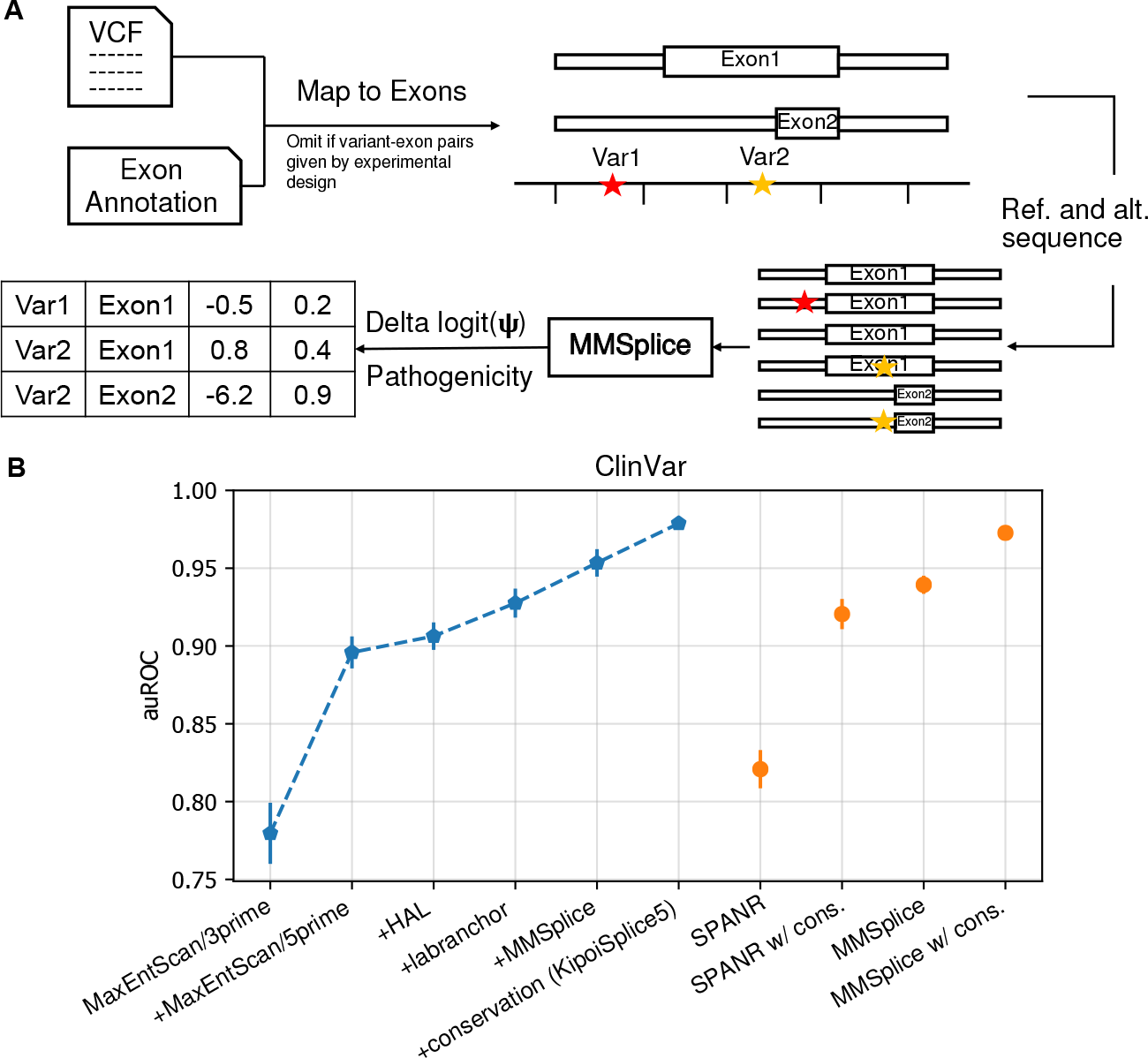
Predictions on ClinVar variants. **(A)** Variants are first mapped to potentially affected exons. Variants from the exon body, within *L_a_* nt from intron of the acceptor sidte or within *L_d_* nt from the donor site are considered to affect splicing of the exon. Afterwards, reference and alternative sequences are retrieved and subjected to MMSplice for prediction. MMSplice gives a prediction for each variant-exon pair. **(B)** Model comparison on classifying pathogenicity of ClinVar splice variants. Models were trained and evaluated in 10-fold cross-validation. Error bars indicate one standard deviation calculated across folds. The six leftmost models (blue) are incrementally added to the ensemble model: ‘+conservation (KipoiSplice5)’ uses all 5 previous models and conservation scores. Performance of MMSplice and SPANR alone as well as their performance with conservation scores are on the right (orange).

This MMSplice model was benchmarked against SPANR [17] and the ensemble of three other models: MaxEntScan [7], HAL [18] and the branch point predictor LaBranchoR [35]. We also compared our MMSplice model and competing models with conservation scores as additional features (Supplementary Methods). Model performances were benchmarked under 10-fold cross-validation (Fig. 5B). MMSplice alone (auROC=0.940) outperformed SPANR (auROC=0.821) (*P* = 0.001, bootstrap) and the ensemble model combining MaxEntScan, HAL and Labranchor (auROC=0.928) (*P* = 0.001, bootstrap). Adding MMSplice to the ensemble model further improved the auROC to 0.954 (*P* = 0.001, bootstrap). Moreover, MMSplice with conservation features (auROC=0.973) achieved a performance close to the best ensemble model kipoiSplice5 that included MMSplice (auROC=0.979) (*P* = 0.003, bootstrap, Fig. 5), indicating that MMSplice alone captured most of the sequence information captured by all other models.

Recently, SPiCE [16] has been proposed as a method to predict the probability of a splice site variant affecting splicing. SPiCE is a logistic regression model trained from 142 manually collected and experimentally tested variants. We thus benchmarked against SPiCE with 12,625 ClinVar variants (2,312 indels) that SPiCE was able to score (it failed to score variants from sex chromosomes)(Methods). MMSplice (auROC = 0.911) outperformed SPiCE (auROC = 0.756 *P* = 0.001, bootstrap). Moreover, this higher performance of the MMPSlice model also held when we fine-tuned the logistic regression model of SPiCE on the ClinVar training dataset (auROC = 0.760 *P* = 0.001, bootstrap, Supplement Fig. S7).

Altogether, these results show that MMSplice not only improves the predictions of effects of variants on biophysical splicing quantities, but also helps improving variant pathogenicity scores.

## Discussion

We have introduced MMSplice, a modular framework to predict the effects of genetic variants on splicing quantities. We did so by training individual modules scoring exon, intron, and splice sites. Models built by integrating these modules showed improved performance against state-of-the-art models on predicting the effects of genetic variants on Ψ, Ψ_3_, Ψ_5_, splicing efficiency, and pathogenicity. The MMSplice software is open source and can be directly applied on VCF files and handles single nucleotide variants and indels. Like other recent models [17, 18, 19], MMSplice score variants beyond the narrow region close to splice sites that is for now suggested by clinical guidelines [36]. We also implemented a VEP [34] plugin that wraps the python implementation. These features should facilitate the integration of MMSplice into bioinformatics pipelines at use in genetic diagnostic centers and may help improving the discovery of pathogenic variants.

MMSplice leverages the modularity of neural networks and deep learning frameworks. MMSplice is implemented using the deep learning python library Keras [37]. All MMSplice modules and models are shared in the model repository Kipoi [26], which should allow other computational biologists to improve individual modules or to flexibly include modules into their own models. We hope this modular approach will help the community to coordinate efforts and continuously and effectively built better variant effect prediction models for splicing.

We have leveraged a massively parallel reporter assay [18] to build individual modules. Also, models predicting Ψ and splicing efficiencies were trained on large-scale perturbation datasets (Vex-seq[24] and MaPSy). Sequence-based model trained from perturbation assays can learn causal features. In contrast, predictive models trained on variations across the reference genome or across natural genetic variations in the population may be limited by evolutionary confounding factors, limiting the model’s ability to make causal predictions about genetic variants. Consistently, our models outperformed models based on the reference genome and natural variations and was only matched by models based on perturbation assays (HAL for ΔΨ_3_ and the SMS score for *in vivo* splicing efficiency changes).

Our models have some limitations. First, splicing is known to be tissue-specific [38, 39], while our models are not. Nevertheless, our models can serve as a good foundation to train tissue-specific models. Second, RNA stability also plays a role in determining the ratio of different isoforms [24]. Models predicting RNA stability from sequence, as we recently developed for the *S. cerevisae* genome [40] could be integrated as further modules. Third, our exon and intron modules are developed from minigene studies, and the performance evaluation on predicting ΔΨ and splicing efficiency changes are also done with minigene experiment data. However, chromatin states is known to has a significant role in splicing regulation [41]. Hence, variant effect prediction for endogenous genes might be benefit from models taking chromatin states into account.

## Methods

### Donor and acceptor modules

The donor and the acceptor modules were trained using the same approach. A classifier was trained to classify positive donor sites (annotated) against negative ones (random, see below) and the same for the acceptor sites. The classifiers predicted scores can be interpreted as predicted strength of the splice sites.

#### Donor and acceptor module training data

For the positive set, we took all annotated splice junctions based on the GENCODE annotation version 24 (GRCh38.p5). For the donor module, a sequence window with 5 nt in the exon and 13 nt in the intron around the donor sites was selected. For the acceptor module, the region around the acceptor sites spanning from 50 nt in the intron to 3 nt in the exon was selected in order to cover most branch points. In total, there were 273,661 unique annotated donor sites and 271,405 unique annotated acceptor sites. This set of splice sites was considered as the positive set. In particular, not only sites with the canonical splicing dinucleotides GT and AG for donor and acceptor sites, respectively, were selected, but also sites with non-canonical splicing dinucleotides were included as positive splice sites.

The negative set consisted of genomic sequences selected within the genes that contributed to positive splice sites, in order to approximately match the sequence context of the positive set. Negative splice sites were selected randomly around but not overlapping the positive splice sites. To increase the robustness of the classifiers, around 50% of the negative splice sites were selected to have the canonical splicing dinucleotides. In total, 410,111 negative donor sites and 406,841 negative acceptor sites were selected. During model training, we split 80% of the data for training and 20% of the data for validation. The best performing model on the validation set was used for variant effect prediction.

#### Donor and acceptor module architecture

Neural network models were trained to score splice sites from one-hot-encoded input sequence. The donor model was a multilayer perceptron with two hidden layers with Rectified Linear Unit (ReLU) activations and a sigmoid output (Supplement Fig.S8A). The hidden layers were trained with a dropout rate [42] of 0.2 and batch normalization [43]. The acceptor model was a convolutional neural network with two consecutive convolution layers (Supplement Fig.S8B). The second convolutional layer was trained with a dropout rate of 0.2 and batch normalization. For these models, we found the number of layers and the number of neurons in each layer by hyperparamater optimization.

### Exon module

#### Exon module training data

The exonic random sequences from the MPRA experiment by [18] were used to train the exon scoring module. This MPRA experiment contains two libraries, one for alternative 5’splicing and one for alternative 3’ splicing. The alternative 5’ splicing library has 265,137 random constructs while the alternative 3’ splicing library has 2,211,789. Each random construct has a 25-nt random sequence in the alternative exon and a 25-nt random sequence in the adjacent intron. Ψ_5_ and Ψ_3_ of different isoforms were quantified by RNA-Seq for each random construct [18]. Here, 80% of the data was used for model training and the remaining were used for validation. The best performing model on the validation set was used for variant effect prediction.

#### Exon module architecture

Rosenberg et al [18] showed that the effects of splicing-related features in alternative exons are strongly correlated with each other across the two MPRA libraries, reflecting that similar exonic regulatory elements are involved for both donor and acceptor splicing. We thus decided to train exon scoring module from the two MPRA libraries by sharing low level convolution layers (Supplement Fig.S1). The inputs of the network were one-hot-encoded 25-nt random sequences. The output labels were Ψ_5_, respectively Ψ_3_, for the alternative exon. After training, the exon modules for each library were separated by transferring the corresponding weights to two separated modules with convolution layer with ReLU non-linearity followed by a global average pooling and a fully connected layer. We have used a global pooling after the convolution layer allowing to take exons of any length as input. This ended up with two exon scoring modules, one for alternative 5’end (exon 5’module) and one for alternative 3’end (exon 3’module).

### Intron module

Intron modules were trained in the same way as the exon modules (Fig. S1) by using intronic random sequences from the MPRA experiment as inputs, except that we used 256 convolution filters, because intronic splicing regulatory elements from the donor side and the acceptor side are less similar [18]. This ended up with a module to score intron on the donor side (intron 5’module) and a module to score intron on the acceptor side (intron 3’module).

### Training procedure for the modules

All neural network models for the six modules were trained with binary cross-entropy loss (Eq.1) and Adam optimizer [44]. We implemented and trained these models with the deep learning python library Keras [37]. Bayesian optimization implemented in hyperopt package [45] was used for hyper-parameter optimization together with the kopt package (github.com/avsecz/kopt). Every trial, a different hyper-parameter combination is proposed by the Bayesian optimizer, with which a model is trained on the training set, its performance is monitored by the validation loss. The model that had the smallest validation loss was selected.

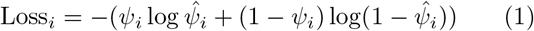

### Variant effect prediction models

#### Variant processing

Variants are considered to affect the splicing of an exon if it is exonic or if it is intronic and at a distance less than *L_a_* from an acceptor site or less than *L_d_* from a donor site. The distances *L_a_* and *L_d_* were set to 100 nt in this study but can be flexibly set for MMSplice. MMSplice provides code to generate reference and alternative sequences from a variant-exon pair by substituting variants into the reference genome. Variant-exon pairs can be directly provided to MMSplice. This is the case for the perturbation assay data Vex-seq, MFASS and MaPSy. MMSplice can also generate variant-exon pairs from given VCF files (Fig. 5A). For insertions, and for deletions that are not overlapping a splice site, the alternative sequence is obtained by inserting or deleting sequence correspondingly. For deletions overlapping a splice site, the alternative sequence is obtained by deleting the sequence and the new splice site is defined as the boundaries of the deletion. In all cases, the returned alternative sequence always have the same structure as the reference sequence, with an exon of flexible length flanked by *L_a_* and *L_d_* intronic nucleotides. Each variant is processed independently from the other variants, i.e. each mutated sequence contains only one variant (Fig. 5A). If a variant can affect multiple target (i.e. sites or exons), the MMSplice models return predictions for every possible target (Fig. 5A).

#### Variant effect prediction for Ψ

Strand information of all Vex-seq assayed exons were first determined by overlapping them with Ensembl GRCh37 annotation release 75. Reference sequences were extracted by taking the whole exon, and 100 nt flanking intronic sequence. Variant sequences were retrieved as described in the variant processing Methods section, whereby variant-exon pairs were provided by the experimental design.

We modeled the differential effect on Ψ in the logistic scale with the following linear model:

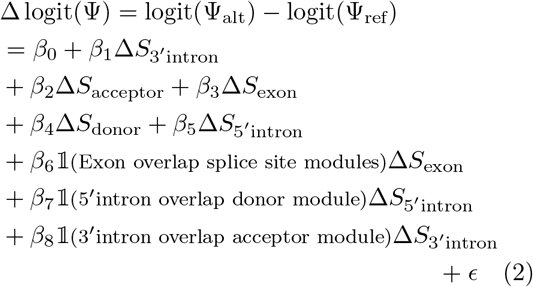

where:

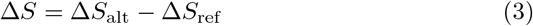

for all five modules, 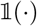 is the indicator function, *ϵ* is the error term, the suffix *alt* denotes the alternate allele, and the suffix *ref* denotes the reference allele. This model has 9 parameters: one intercept, one coefficient for each of the five modules, and interaction terms for regions that were scored by two modules (Fig.1). The latter interaction terms were useful to not double count the effect of variants scored by multiple modules. These 9 parameters were the only parameters that were trained from the Vex-seq data. The parameters of the modules stayed fixed. To fit this linear model, we used Huber loss [46] instead of ordinary least squares loss to make the fitting more robust to outliers.

The model predicts Δ logit Ψ for the variant. We transform this to ΔΨ with a given reference Ψ as follow:

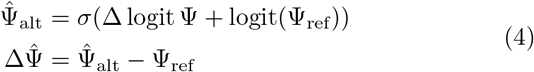

where:

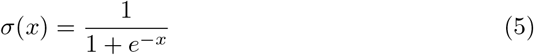

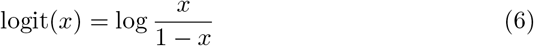

To prevent infinite values in cases Ψ_ref_ = 0 or Ψ_ref_ = 1, Ψ_ref_ values were clipped to the interval [10^−5^, 1−10^−5^].

HAL model is provided by the authors. A scaling factor required by HAL was trained on the Vex-seq training data using code provided by the authors [18]. The SPANR precomputed scores (which are called SPIDEX), were obtained from http://www.openbioinformatics.org/annovar/spidex_download_form.php.

#### Performance on the MFASS dataset

MMSplice was applied the same way as for Vex-seq, except that module combining weights were learned from the Vex-seq training data, with MFASS data kept entirely unseen. SDVs are classified based on the predicted ΔΨ for a variant. Area under the precision-recall curve (auPR) were calculated with `trapz` function from R package `pracma`.

#### Variant effect prediction for Ψ_3_ and Ψ_5_

The Genotype-Tissue Expression (GTEx) [32] RNAseq data (V6) was used to extract variant effect on Ψ_3_ and Ψ_5_. Variants [−3, +6] nt around alternative donors of alternative 5’ splicing events and variants [−20, +3] nt around alternative acceptors for alternative 5’ splicing events were considered. The skin (not sun exposed) samples and the brain samples with matched whole genome sequence data available were processed. Ψ_5_ and Ψ_3_ were calculated with MISO [20] for each sample. Altogether, 1,057 brain samples and 211 skin samples could be successfully processed with MISO. Ψ_3_ and Ψ_5_ for homozygous reference variant, heterozygous variants, and homozygous alternative variants were calculated by taking the average across samples with the same genotype, excluding samples from individuals with more than one variants within 300 nt around the competing splice sites.

We predicted differences in Ψ_5_ as follows. We considered only donor sites with two alternative acceptor sites. We extracted the relevant sequences for the corresponding two alternative exons and apply the model of Equation (2) which was fitted on Vex-seq training data. This returned a Δlogit(Ψ) for each alternative exon, denoted Δ*S*_1_ and Δ*S*_2_, from which we calculate the predicted alternative Ψ_5_ as follows (derivations provided in supplements):

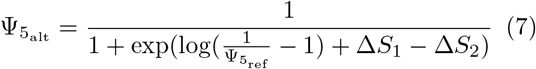

The above computation applies to individual alleles. To handle heterozygous variants, we assumed expression from both alleles are equal. This led to the following predictions for homozygous and heterozygous variants:

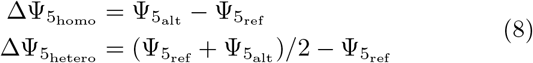

Analagous calculations were made to predict differences in Ψ_3_.

Pre-trained COSSMO model [19] was obtained from the author website (http://cossmo.genes.toronto.edu/). The predicted ΔΨ_5_ (or ΔΨ_3_) values of COSSMO were calculated by taking the difference between the predicted Ψ_5_ (or Ψ_3_) from alternative sequence processed by MMSplice and reference sequence.

#### Splicing efficiency dataset (MaPSy data)

The splicing efficiency assay was performed for 5,761 disease causing exonic nonsynonymous variants both *in vivo* in HEK293 cells and *in vitro* in HeLa-S3 nuclear extract as previously described [22]. Here, the exons were derived from human exons and were reduced in size to be shorter than 100 nt long by small deletions applied to both the reference and the alternative version of the sequence. This way, the wild-type and the mutated alleles differed from each other by a single point mutation and the wild-type allele differed from a human exon by the small deletions. The deletions were centered at the midpoint between the variant and the furthest exon boundary. The sequences of each substrate are listed in Supplemental Table 1 and also described further on the CAGI website (https://genomeinterpretation.org/content/MaPSy).

Overall, 4,964 of the variants were in the training set and 797 were in the test set. The amount of spliced transcripts and unspliced transcripts for each construct with reference allele or alternative allele were determined by RNA-Seq. The effect of mutation on splicing efficiency for a specific reporter sequence was quantified by the allelic log-ratio, which is defined as:

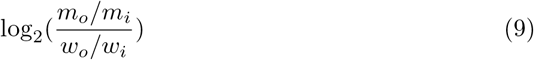

where *m_o_* is the mutant spliced RNA read count, *m_i_* is the mutant input (unspliced) RNA read count, *w_o_* is the wild-type spliced RNA read count, *w_i_* is the wild-type input RNA read count. Transcripts with exon-skipped or misspliced are ignored.

#### Variant effect prediction for splicing efficiency (MaPSy data)

We fitted a model to predict differential splicing efficiency on the training set with a linear regression with a Huber loss as defined by Equation 2, except that the response variable is the allelic log-ratio (Equation 9) instead of Δ logit(Ψ). We used the exon 5’module for the splicing efficiency model. Performance on MaPSy data was reported on the held-out test set.

SMS scores was applied to wild-type and mutant sequence by summing up all 7-mer scores as described by Ke et al. [23]. The predicted allelic log-ratio is the SMS score difference between mutant and wil-type sequence.

#### Variant pathogenicity prediction

Processed ClinVar variants (version 20180429 for GRCh37) around splcie sites were obtained from Avsec et al. [26]. Specifically, single-nucleotide variants [−40, 10] nt around the splicing acceptor or [−10, 10] nt around the splice donor of a protein coding gene (Ensembl GRCh37 v75 annotation) were selected. Variants causing a premature stop codon were discarded. After the filtering, the 6,310 pathogenic variants constituted the positive set and the 4,405 benign variants constituted the negative set. The CADD [30] scores and the phyloP [47] scores were obtained through VEP [34]. MMSplice Δ*Score* predictions of the 5 modules as well as indicator variables of the overlapping region were assembled with a logistic regression model to classify pathogenicity. Performance was assessed by 10-fold cross-validation (Supplementary Methods).

To compare MMSplice with SPiCE [16], we restricted to the regions that SPiCE scores, i.e. [−12, 2] nt around the acceptor or [−3, 8] nt around the donor of protein coding genes. Variants causing a premature stop codon were discarded. SPiCE was trained to predict the probability of a variant to affect splicing (manually defined by experimental observations). To apply it for pathogenicity prediction, the logistic regression model of SPiCE was refitted with ClinVar pathogenicity as response variable. MMSplice model was applied as described above without conservation features. Models were compared under 10-fold cross-validation.

#### P-values for model performance comparison

Significance levels when comparing the performance of two models were estimated with the basic bootstrap [48]. Denoting *t*_1_ the performance metric (Pearson correlation, auPRC, or auROC) of MMSplice and *t*_2_ the performance metric of a competing model, we considered the difference *d* = *t*_1_ − *t*_2_. We sampled with replacement the test data *B* = 999 times and each time *i* computed the bootstrapped metric difference 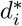. The one-sided P-value was approximated as [48].

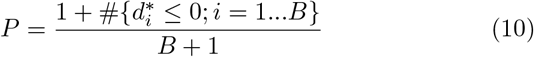

## Acknowledgements

We thank the Critical Assessment of Genome Interpretation (CAGI) organizers, especially Steven E. Brenner, John Moult and Gaia Andreoletti for organizing the CAGI competition. We thank Scott I. Adamson, Brenton R. Graveley for formatting and providing Vex-seq experiment data through CAGI.

## Availability of data and materials

MMSplice and its VEP plugin is available from https://github.com/gagneurlab/MMSplice and Kipoi: https://kipoi.org/models/MMSplice. Analysis code is available from https://github.com/gagneurlab/MMSplice_paper. Vex-seq data is available from https://github.com/scottiadamson/Vex-seq. MFASS data is available from https://github.com/KosuriLab/MFASS. GTEx data is available from dbGAP (phs000424.v6.p1) https://www.ncbi.nlm.nih.gov/projects/gap/cgi-bin/study.cgi?study_id=phs000424.v6.p1. ftp://ftp.ncbi.nlm.nih.gov/pub/clinvar/vcf_GRCh37/archive_2.0/2018/clinvar_20180429.vcf.gz. MaPSy data is available in the Additional files for this manuscript.

## Author contributions

JC and JG designed the model, with the help of ZA. JC implemented the software and analysed data. TYDN, ZA contributed to developing the modules. JC and JG wrote the manuscript, with the help of ZA, KJC and WGF. KC and WF generated the MaPSy data. MHC wrote the VEP plugin.

